# *At*KC1 inhibits *At*AKT1 activation via its amino-terminal inhibitory domain

**DOI:** 10.1101/2024.12.27.630498

**Authors:** Zijie Zheng, Yannan Qu, Jiexin Chen, Yuyue Tang, Dongliang Liu, Zhuo Huang, Huaizong Shen

## Abstract

AKT1 is a plant Shaker-like, hyperpolarization-activated K^+^ channel which plays a crucial role in K^+^ absorption. Besides phosphorylation, AKT1 is subject to negative regulation by *At*KC1, a silent channel of the same family. Previous structural studies unveiled that AKT1 and *At*KC1 form 2:2 heterotetramer in purified samples. However, the structural analysis failed to offer more insight into the inhibition mechanism of AKT1 activation by *At*KC1. Here, inspecting the complex structure of AKT1-*At*KC1 reveals that a stable domain of *At*KC1 (residues 53 to 80), named inhibitory domain or I-domain, may inhibit AKT1 activation by stabilizing its depolarized, closed conformation. We confirmed this hypothesis with electrophysiological experiments. Interestingly, a single-point mutation (G315D) in *At*KC1 has been reported to abolish its ability to inhibit AKT1. We solved the structure of AKT1-*At*KC1(G315D) at a resolution of 2.8 Å which revealed an unexpected stoichiometry alteration between AKT1 and *At*KC1 from 2:2 to 3:1. This stoichiometry change further supports the hypothesis as we reason that single I-domain of *At*KC1 in the channel complex is insufficient to effectively inhibit channel activation. Our findings reveal the inhibition mechanism of *At*KC1 on AKT1 and offer insight into the regulatory mechanisms of hyperpolarization-activated channels.

## Introduction

Potassium (K^+^) is an essential macronutrient for the growth and development of plants. It performs various vital functions within plant cells, including osmoregulation, anion neutralization, enzyme activation, and membrane transport processes^1–7^. In *Arabidopsis thaliana*, there are two major high-affinity K^+^ transport systems, AKT1 (<u>A</u>rabidopsis <u>K</u>^+^ transporter <u>1</u>) and HAK5 (<u>h</u>igh-<u>a</u>ffinity <u>K</u>^+^ transporter <u>5</u>), responsible for the uptake of K^+^ from soils^6,8–12^.

AKT1 was initially identified through a functional complementation assay in yeast cells deficient in K^+^ absorption^13^. Sequence analysis and functional characterization revealed that AKT1 is a Shaker-like, hyperpolarization-activated K^+^ channel^4,13^. It consists of the intramembrane voltage-sensing domain (VSD), pore domain (PD), and cytoplasmic domains, including the C-linker, cyclic nucleotide-binding homology domain (CNBHD), ankyrin repeat domain (ANK domain), and K_HA_ domain (Supplementary Fig. S1)^4,13^. Genetic and functional studies have demonstrated that the activity of AKT1 is tightly regulated by a calcium-related kinase-phosphatase system through phosphorylation or dephosphorylation of residues within AKT1’s cytoplasmic domains, with Calcineurin B-like protein 1 or 9 (CBL1/9) and CBL-interacting serine/threonine-protein kinase 23 (CIPK23) required for the activation of AKT1^14–18^. Additionally, AKT1 is inhibited by a homolog, *At*KC1, which is crucial for plant tolerance under low-K^+^ conditions^19–22^. *At*KC1 has been proved a silent channel that functions by regulating the properties of homologous channels such as AKT1 and KAT1, potentially through heterotetramer formation^23–27^. Interestingly, a single-point mutation (G315D) in *At*KC1 has been reported to completely abolish its ability to inhibit AKT1 activation through unknown mechanisms^20^.

Recent studies have reported the structures of AKT1 and the AKT1-*At*KC1 complex^28,29^. The AKT1-*At*KC1 complex structure reveals the formation of a heterotetramer with a 2:2 ratio between AKT1 and *At*KC1. Surprisingly, both structures of AKT1 and the AKT1-*At*KC1 complex exhibit a C2 symmetry in the cytoplasmic domains rather than a C4 symmetry. Based on structural and mutagenesis studies, it is proposed that the transition of the cytoplasmic domains from C2 to C4 symmetries is involved in the activation process of AKT1 and AKT1-*At*KC1. However, further evidence is required to confirm this hypothesis.

Although the AKT1-*At*KC1 structure revealed the heterotetramerization of the two proteins in the channel complex, previous structural analysis failed to provide more insight into the inhibition mechanism of AKT1 by *At*KC1^28,29^. In this study, a combination of structural and electrophysiological methods was employed to specifically investigate the inhibition mechanism of AKT1 activation by *At*KC1.

## Results

### Electrophysiological characterization and structural determination of AKT1 and AKT1-*At*KC1

The RNAs encoding AKT1, CBL1, and CIPK23 were transcribed in vitro and subsequently transfected into oocytes, either with or without the RNA for *At*KC1. The transfected oocytes were then subjected to electrophysiological examination using two-electrode voltage clamps (TEVC) in bath solutions containing 96- or 0.5-mM K^+^ ions (Fig. 1a and Supplementary Fig. S2). Consistent with previous studies, our electrophysiological data demonstrate that AKT1 functions as a hyperpolarization-activated K^+^ channel that can be effectively inhibited by *At*KC1 at hyperpolarized membrane potentials^20–22,29^.

**Figure 1.**
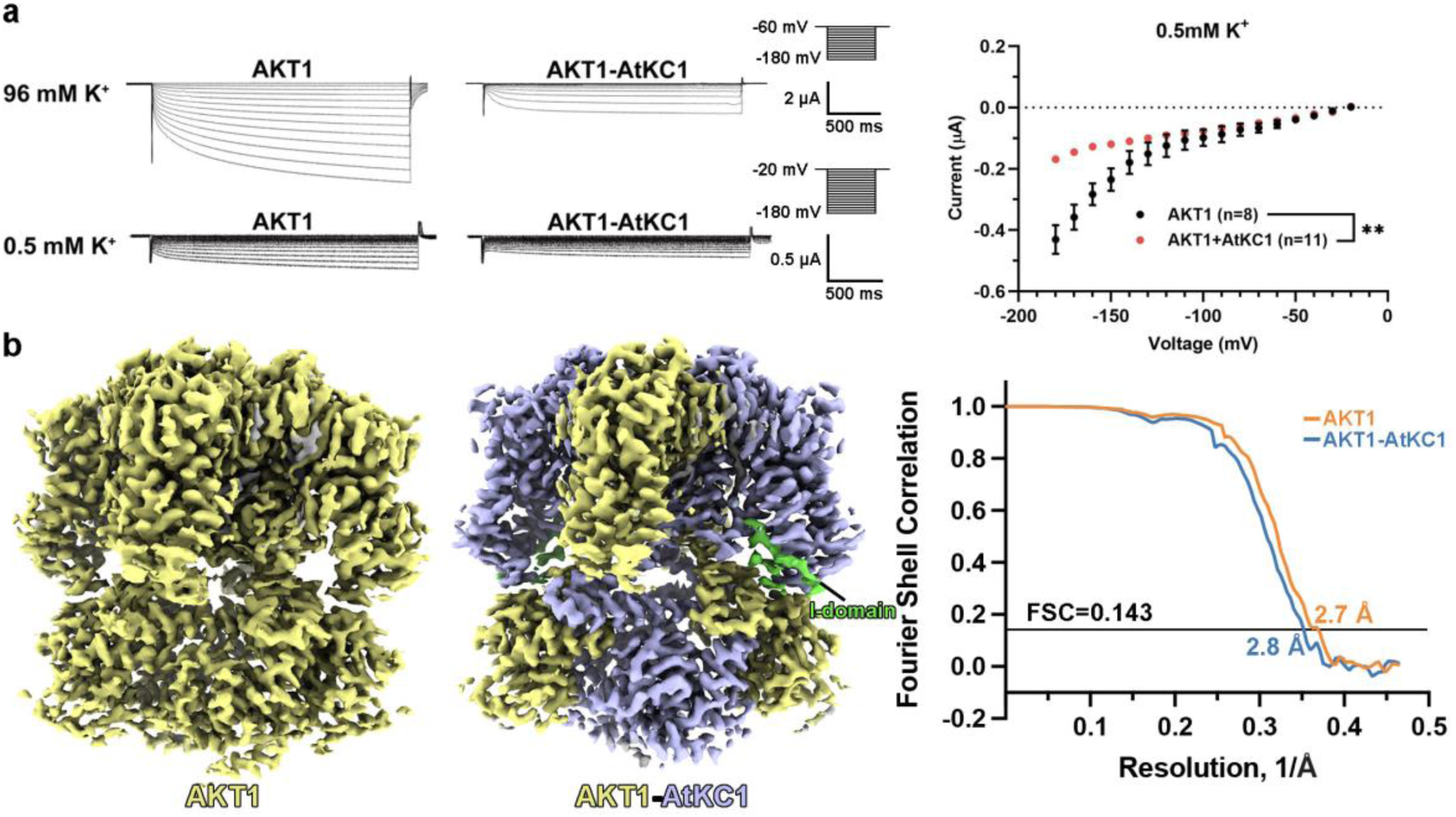
Characterization and structural determination of AKT1 and AKT1-*At*KC1. (a) Electrophysiological characterization of AKT1 and AKT1-*At*KC1 in oocytes. AKT1 or AKT1-*At*KC1 is co-transfected with CBL1 and CIPK23 in oocytes, which were electrophysiologically examined by two-electrode voltage clamps at indicated series of voltages. Representative traces (left) and corresponding I-V curves (right) of AKT1 and AKT1-*At*KC1 were recorded in bath solutions containing indicated concentrations of K^+^. Comparisons among groups were made using two-way analysis of variance followed by Bonferroni’s post hoc test. * p < 0.05, ** p < 0.01, *** p < 0.001. All data are presented as mean ± SEM. (**b**) Structural determination of AKT1 and AKT1-*At*KC1. AKT1 forms a homotetramer, whereas AKT1-*At*KC1 exists as a heterotetramer with a 2:2 ratio of AKT1 and *At*KC1. Fourier shell correlation (FSC) curves indicate resolutions of 2.7 and 2.8 Å for the reconstructions of AKT1 and AKT1-*At*KC1, respectively, as determined by the gold-standard FSC 0.143 criterion^60^. EM map figures were prepared using UCSF ChimeraX^66^.

Following standard protocols, the cryo-EM structures of AKT1 and AKT1-*At*KC1 were determined at resolutions of 2.7 and 2.8 Å, respectively (Fig. 1b, Supplementary Figs. S3-S6, and Table S1). The cryo-EM maps resolve the densities for the transmembrane domain, cytoplasmic C-linker, and CNBHD domain of both AKT1 (residues 52 to 492) and *At*KC1 (residues 53 to 526). The structures closely resemble those independently reported by two other research groups and exhibit similar features (Supplementary Fig. S7)^28,29^. Both AKT1 and *At*KC1 adopt the non-domain-swapped configuration due to the presence of short S4-S5 linkers (∼6 residues)(Fig. 2a and Supplementary Fig. S8), which is also observed in the structures of human K_v_10-12, CNG, HCN channels, and KAT1 from Arabidopsis^30–35^. Furthermore, both proteins display depolarized VSDs and closed pores, similar to the structure of KAT1 (Supplementary Fig. S9)^34,35^. Additionally, a symmetry reduction from C4 to C2 for the cytoplasmic domains is also observed in our structures, confirming previous structural findings (Fig. 2a and Supplementary Fig. S8)^28,29^.

**Fig. 2.**
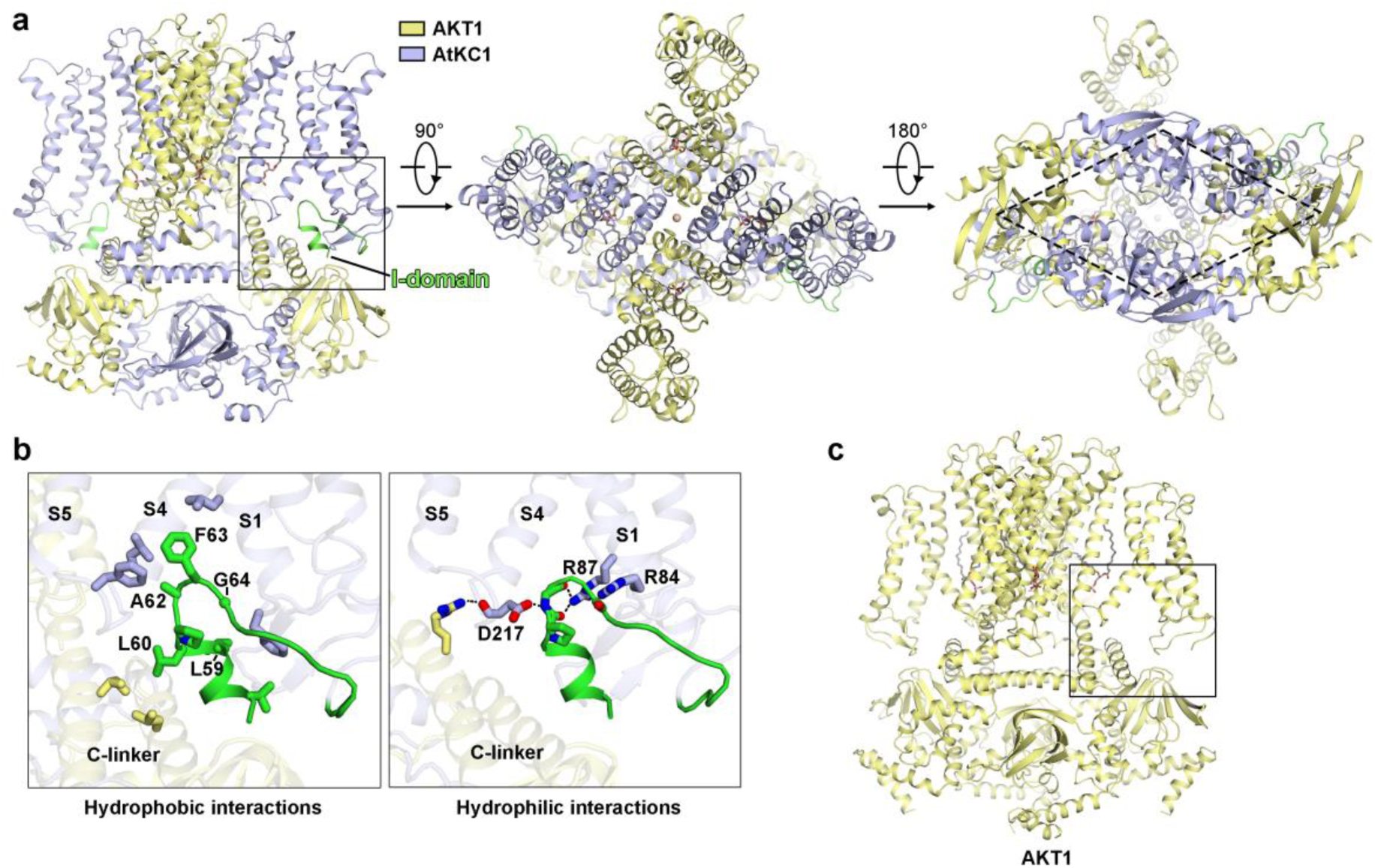
Extensive interactions of the I-domain in *At*KC1 with surrounding structural elements. (**a**) Overall structure of AKT1-*At*KC1 from different viewing angles. The cytoplasmic domain of AKT1-*At*KC1 adopts a rhombus-like shape of a rhombus with C2 symmetry instead of C4 symmetry. The I-domain of *At*KC1 (green) inserts into the cavity formed by the S1, S4, and S4-S5 linker of the same polypeptide chain and the C-linker of an adjacent AKT1. (**b**) The I-domain of *At*KC1 engages in extensive hydrophobic and hydrophilic interactions with adjacent structural elements. Protein segments and residues involved in the interactions are labeled. (**c**) In contrast to the presence of the I-domain of *At*KC1 in the AKT1-*At*KC1 structure, the corresponding sequence of AKT1 is absent in both AKT1 and AKT1-*At*KC1 structures, suggesting its flexibility. All structure figures were prepared using PyMOL^67^.

### Inspection of AKT1-*At*KC1 reveals a potential inhibitory domain of *At*KC1

Comparison of the structures of AKT1 and AKT1-*At*KC1 reveals a notable difference: a stable domain of *At*KC1 (residues 53 to 80) preceding the transmembrane domain occupies the cavity enclosed by the S1, S4, S4-S5 linker from the same subunit, and the C-linker from an adjacent AKT1 subunit (Fig. 2a and Supplementary Fig. S6). Subsequent studies indicate that this domain of *At*KC1 may play an inhibitory role in AKT1 activation, thus we named this domain (residues 53 to 80) the inhibitory domain or the I-domain. In contrast, the corresponding regions in AKT1 are invisible in both structures (Fig. 2c and Supplementary Fig. S8).

The I-domain of *At*KC1 forms extensive interactions, both hydrophobic and polar ones, with its surrounding structural elements (Fig. 2b). Firstly, the hydrophobic loop (_59_LLPAFG_64_) of the I-domain strongly favors docking into the cytoplasm-facing cavity surrounded by hydrophobic residues from S1, S4 and S4-S5 linker of the same subunit, as well as the C-linker of an adjacent AKT1 subunit, with the _62_AFG_64_ residues inserting most deeply. Notably, a conserved Phe residue in HCN channels is present at a similar location as Phe63 in *At*KC1^32,36^. Secondly, the domain is further stabilized by local polar interactions between the main-chain nitrogen atoms, carbonyl groups, and the charged side chains of the aforementioned segments. Thirdly, the I-domain of *At*KC1 also interacts with the S2-S3 linker of the same subunit. We hypothesized that these extensive interactions between the I-domain of *At*KC1 and the surrounding structural elements are responsible for the inhibition mechanism of AKT1 activation by *At*KC1 through stabilizing the depolarized, closed conformation of the channel complex present in the solved structure (Supplementary Fig. S9).

### Electrophysiological assays on AKT1 and *At*KC1 mutants

To validate our hypothesis regarding the inhibitory role of the I-domain of *At*KC1 in the activation of AKT1, we designed several AKT1 and *At*KC1 mutants based on sequence alignment between AKT1 and *At*KC1 (Fig. 3 and Supplementary Fig. S1). Firstly, we replaced the N-terminal region of AKT1 (residues 1-51) with the N-terminal sequence (residues 1-80, including the I-domain) of *At*KC1, and named the resulting mutant AKT1-I. In comparison with the wild-type AKT1, AKT1-I exhibits substantially smaller currents at hyperpolarized membrane potentials, similar to those recorded for AKT1-*At*KC1 (Fig. 3a). Furthermore, when we mutated the three hydrophobic residues (_62_AFG_64_), which are deeply inserted into the docking site of the I-domain, to three consecutive hydrophilic glutamine residues (QQQ), the inhibitory effect exerted by the I-domain on AKT1-I was completely abolished. Consistent results were obtained when *At*KC1 was electrophysiologically assayed after removing its N-terminus (*At*KC1-ΔI) or replacing the _62_AFG_64_ residues with QQQ (*At*KC1-QQQ) (Fig. 3b). Both AKT1-*At*KC1-ΔI and AKT1-*At*KC1-QQQ displayed I-V curves that indicated a loss of the inhibitory effects of *At*KC1 on AKT1.

**Fig. 3.**
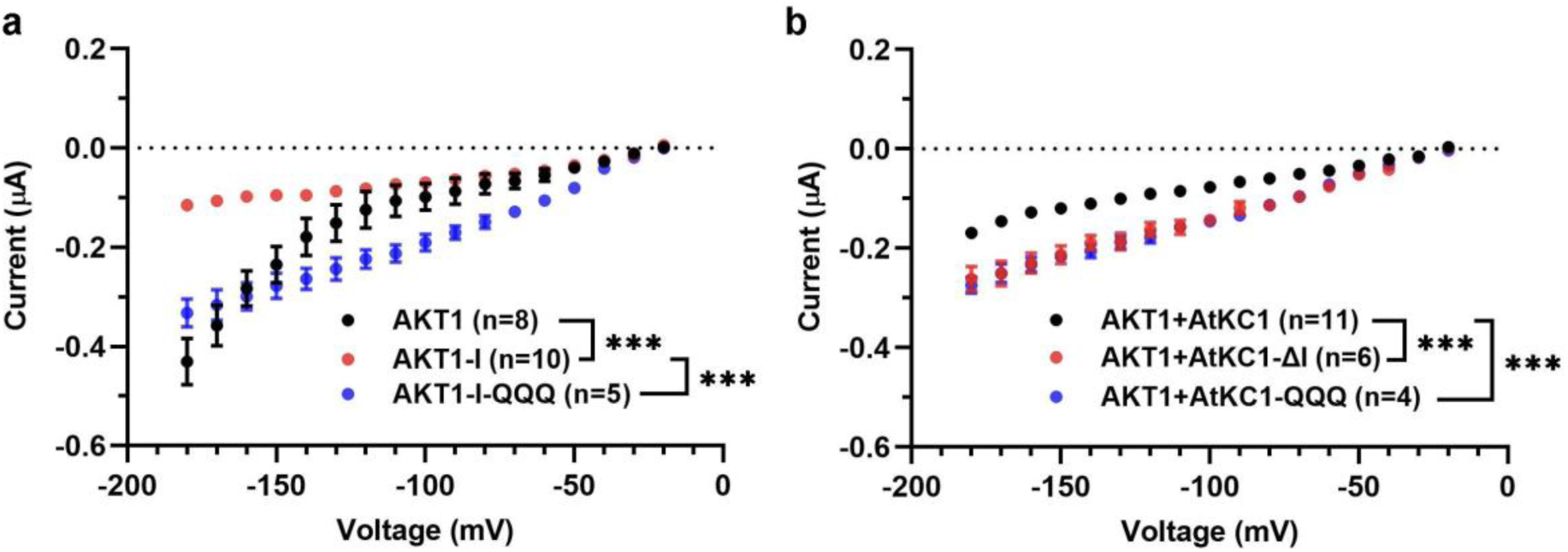
Electrophysiological recordings of AKT1, *At*KC1, and their mutants. (**a**) I-V curves of AKT1, AKT1-I (AKT1 with its N-terminus (residues 1-51) replaced by the corresponding region of *At*KC1 (residues 1-80)), and AKT1-I-QQQ (AKT1-I with the _62_AFG_64_ residues mutated to QQQ) were recorded in bath solutions containing 0.5 mM K^+^. (**b**) I-V curves of AKT1-*At*KC1, AKT1-*At*KC1-ΔI (*At*KC1 with the deletion of its N-terminus (residues 1-80)), and AKT1-*At*KC1-QQQ (*At*KC1 with the _62_AFG_64_ residues mutated to QQQ) were recorded in bath solutions containing 0.5 mM K^+^. These I-V curves provide insight into the inhibitory roles of the I domain and _62_AFG_64_ residues of *At*KC1 in the activation of AKT1. Comparisons among groups were made using a two-way analysis of variance followed by Bonferroni’s post hoc test. * p < 0.05, ** p < 0.01, *** p < 0.001. All data are presented as mean ± SEM.

### Stoichiometry alteration between AKT1 and *At*KC1 in AKT1-*At*KC1(G315D)

The G315D mutation in *At*KC1 has previously been reported to abolish its inhibitory effect on AKT1^20^. To gain further insight into the inhibition mechanism of *At*KC1 on AKT1, we determined the cryo-EM structure of AKT1-*At*KC1(G315D) at a resolution of 2.8 Å (Fig. 4a, Supplementary Figs. S3, S4, and Table S1). Unexpectedly, the G315D mutation caused a complete change in the stoichiometry between AKT1 and *At*KC1 from 2:2 in AKT1-*At*KC1 to 3:1 (Fig. 4). Asp315 of *At*KC1(G315D) is situated in the middle of the transmembrane segment S6, with its carboxyl group in the side chain pointing unfavorably towards the hydrophobic lipid bilayers (Fig. S10). We propose that the energetic penalty imposed by the unfavorable accommodation of an aspartic residue in the lipid environment prevents the simultaneous incorporation of two AtKC1 subunits into one channel complex. As a result, only one I-domain of *At*KC1 is present in the AKT1-*At*KC1(G315D) complex, which may be insufficient to effectively inhibit the activation of AKT1.

**Fig. 4.**
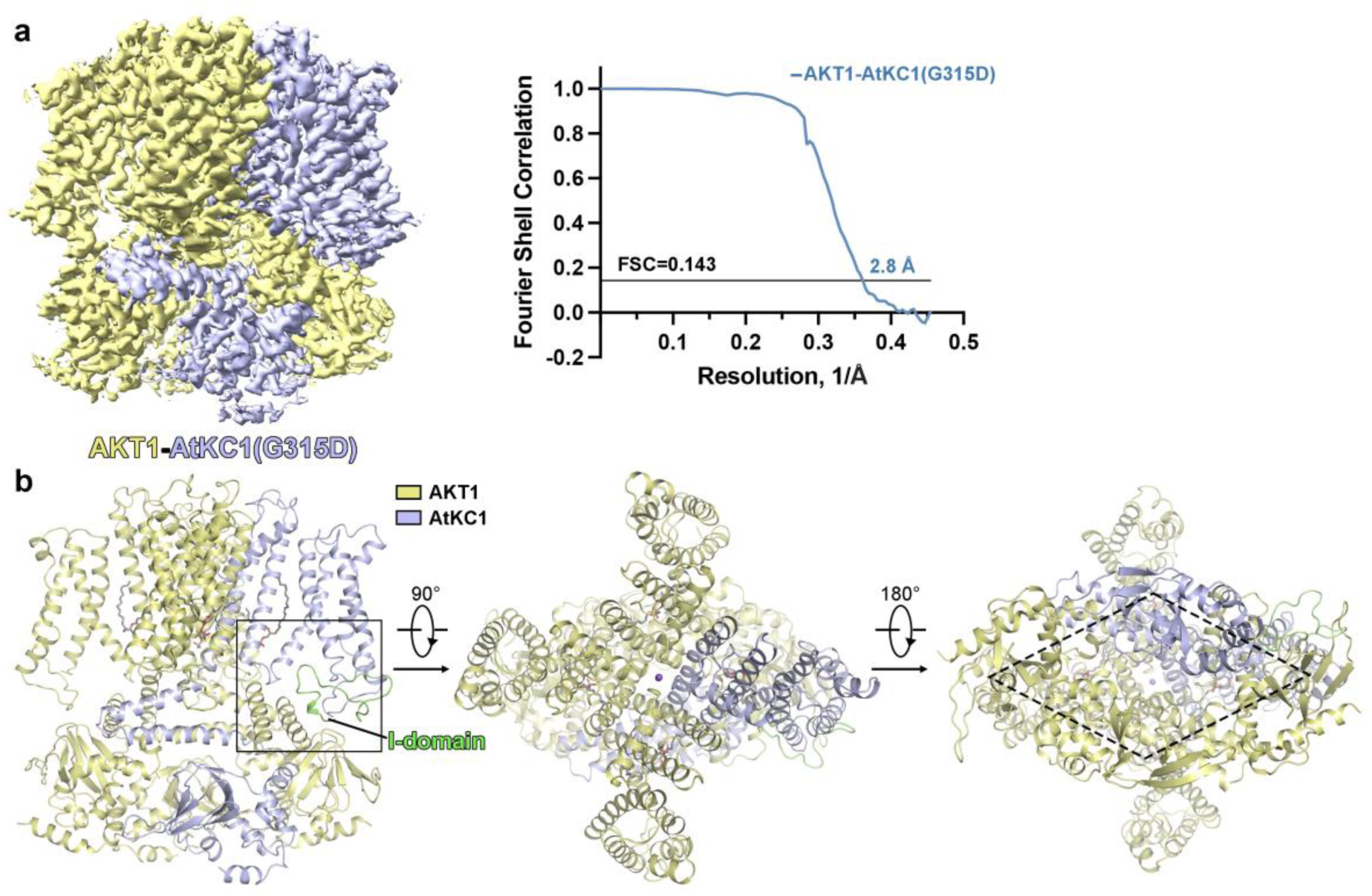
Structure of AKT1-*At*KC1(G315D) reveals a 3:1 ratio between AKT1 and *At*KC1(G315D). (**a**) The cryo-EM structure of AKT1-*At*KC1(G315D) determined at a resolution of 2.8 Å demonstrates an altered stoichiometry between AKT1 and *At*KC1, transitioning from a 2:2 ratio in AKT1-*At*KC1 to a 3:1 ratio in AKT1-*At*KC1(G315D). **(b)** Overall structure of AKT1-*At*KC1(G315D) from different viewing angles. The cytoplasmic domain of AKT1-*At*KC1(G315D) also assumes a rhombus-like shape. The stoichiometry alteration between AKT1 and *At*KC1 results in the presence of only one I-domain of *At*KC1 (green) in AKT1-*At*KC1(G315D).

### Model for the inhibition mechanism of *At*KC1 on AKT1

Based on the structural and electrophysiological observations, we propose a model for the inhibition mechanism of *At*KC1 on AKT1 (Fig. 5). In the depolarized conformation as present in the solved structures, the AKT1-*At*KC1 channel complex is closed, with the I-domain of *At*KC1 interacts extensively with surrounding structural elements, thereby stabilizing the closed conformation. Upon activation by hyperpolarization, the AKT1-*At*KC1 complex undergoes dramatic conformational changes, including the initial downward movement of the S4 segments towards the cytoplasmic side and the subsequent replacement of the S1, S4, S2-S3, S4-S5 linkers, and the cytoplasmic C-linkers. These dramatic conformational changes are presumably hindered by the extensive interactions between the I-domains of *At*KC1 and their surrounding structural elements. Therefore, the release of the I-domains or the disruption of their interactions with surrounding residues should be a prerequisite for the activation of the channel complex. This model explains the mechanism through which *At*KC1 inhibits the activation of AKT1.

**Fig. 5.**
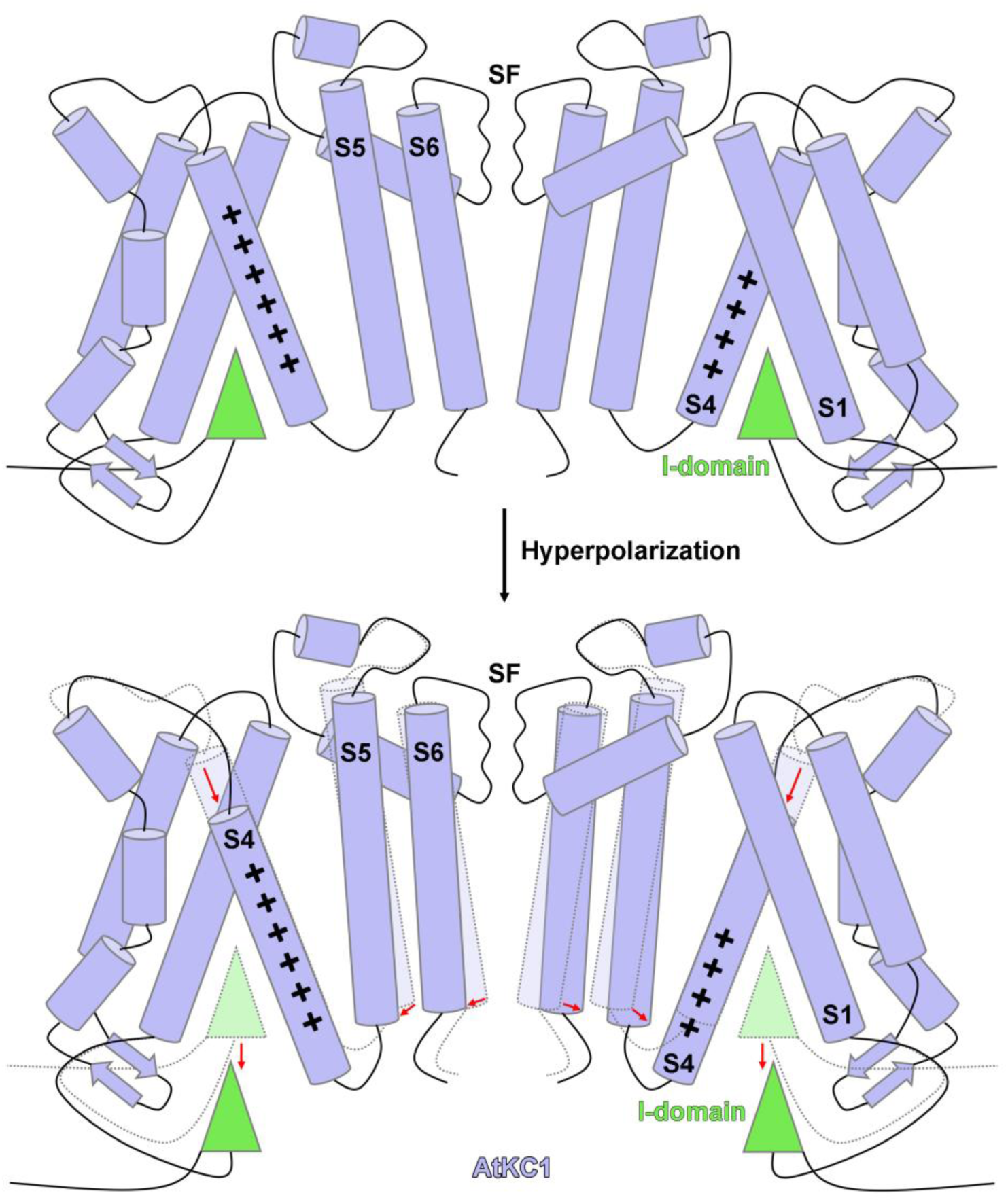
Model for the inhibition mechanism of *At*KC1 on AKT1 activation. Based on our structural and electrophysiological findings, we propose a model for the inhibition mechanism of *At*KC1 on AKT1 activation. Upon membrane potential hyperpolarization-induced activation, the S4 segments of AKT1 and *At*KC1 undergo movement towards the cytoplasmic side, leading to substantial structural changes in the S1, S4-S5 linker, and C-linkers. Consequently, these altered segments can no longer accommodate the stable insertion of the I-domain of *At*KC1. Therefore, these extensive interactions between the I-domain of *At*KC1 and its accommodating site residues in the channel’s closed state play a critical role in inhibiting the activation process of AKT1. The I-domain of *At*KC1 is represented by green triangles, and different segments are appropriately labeled. For clarity, C-linkers are omitted from the model.

## Discussion

AKT1 is a hyperpolarization-activated, CNBHD domain-containing channel, making it a suitable subject for studying the activation, inhibition, and other regulatory mechanisms of channels with similar gating properties or structures^32,33,37–45^.

The discovery of the inhibitory role of the I-domain in *At*KC1 on AKT1 here is reminiscent of the HCN domain in HCN channels, which has also been reported to prevent the channel from activation by stabilizing the closed state^32,36^. The docking site of the I-domain, which is composed of the S4 and other surrounding segments, plays a crucial role in the electromechanical coupling of voltage-gated ion channels (VGICs)^35,36,46–49^. It is not surprising that evolution has devised regulatory mechanisms around this site.

The alteration in the stoichiometry between AKT1 and *At*KC1 in the AKT1-*At*KC1 and AKT1-*At*KC1(G315D) complex structures is unanticipated. To the best of our knowledge, this is the first time that a single-point mutation in the S6 segment of a voltage-gated ion channel (VGIC) has been found to cause a complete alteration in the stoichiometry of the constituting subunits. Similar strategies may be employed to manipulate the subunit compositions of other VGICs.

In summary, our structural and electrophysiological studies unveil that *At*KC1 inhibits AKT1 via its N-terminal inhibitory domain. These findings provide insight into the regulatory mechanisms of other similar channels, such as human K_v_10-12, CNG, and HCN channels. Additionally, considering the crucial roles of AKT1 and *At*KC1 in plant tolerance to low-K^+^ conditions and salinity stress, our results hold great potential in developing new strategies for increasing crop yields.

## Materials and methods

### Electrophysiological recordings

The coding sequences of AKT1, AKT1-I, AKT1-I-QQQ, *At*KC1, *At*KC1-ΔI, *At*KC1-ΔI-QQQ, CBL1, and CIPK23 were cloned into pKSM vector and linearized by Enzyme (SacⅡ). The cRNAs were transcribed in vitro using the T3 RNA polymerase (Thermo, USA). Oocytes were isolated from Xenopus laevis and injected with cRNAs. Stage V and VI oocytes were injected with AKT1, CBL1 and CIPK23 cRNA mixture (3.83:1.92:1.92 ng in 23 nL), AKT1, *At*KC1, CBL1 and CIPK23 cRNA mixture (3.83:3.83:1.92:1.92 ng in 23 nL), AKT1-I, CBL1 and CIPK23 cRNA mixture (3.83:1.92:1.92 ng in 23 nL), AKT1-I-QQQ, CBL1 and CIPK23 cRNA mixture (3.83:1.92:1.92 ng in 23 nL), AKT1, *At*KC1-ΔI, CBL1 and CIPK23 cRNA mixture (3.83:3.83:1.92:1.92 ng in 23 nL), AKT1, *At*KC1-ΔI-QQQ, CBL1 and CIPK23 cRNA mixture (3.83:3.83:1.92:1.92 ng in 23 nL). The injected oocytes were incubated for 1-3 days at 16°C in ND96 solution (96 mM NaCl, 2 mM KCl, 5 mM MgCl2, 1.8 mM CaCl2, 5 mM HEPES, pH 7.4).

Whole-oocyte currents were measured by a two-electrode voltage clamp amplifier OC-725C (Warner Instruments). Electrodes were filled with 3 M KCl and had resistances of 0.1–1.0 MΩ. The high K^+^ bath solution contained (in mM) 96 KCl, 1.8 MgCl_2_, 1.8 CaCl_2_, 0.1 LaCl_3_, 10 HEPES-NaOH (pH 7.2). The low K^+^ bath solution contained (in mM) 0.5 KCl, 1.8 MgCl_2_, 1.8 CaCl_2_, 0.1 LaCl_3_, 10 HEPES-NaOH (pH 7.2). The low K^+^ bath solution contained D-mannitol to adjust the osmolality. For measurements of channel activation in high K^+^ bath solution, oocytes were held at −60 mV, and 2500-millisecond duration test hyperpolarizations were applied in 10 mV increments, from −60 mV to −180 mV, every 2 seconds. For measurements of channel activation in low K^+^ bath solution, oocytes were held at −20 mV, and 2500-millisecond duration test hyperpolarizations were applied in 10 mV increments, from −20 mV to −180 mV, every 2 seconds. Data were filtered at 1 kHz. Data acquisition and analysis were carried out with Digidata 1322A (Axon Instruments Inc., Foster City, CA, USA) and pCLAMP software.

Statistical analyses were performed using Clampfit 10.7 (Molecular Devices, LLC, USA) and GraphPad Prism 7 (GraphPad Software, La Jolla, CA). Comparisons among groups were made using two-way analysis of variance followed by Bonferroni’s post hoc test. * p < 0.05, ** p < 0.01, *** p < 0.001. All data are presented as mean ± SEM.

### Cloning, expression, and purification of AKT1, AKT1-*At*KC1 and AKT1-*At*KC1(G315D)

For the structural study of AKT1, gene encoding *Arabidopsis thaliana* AKT1 (UniProt: Q38998) was synthesized by GENEWIZ and confirmed by sequencing. Full-length coding sequence was cloned into a pCAG expression vector with the DrICE protease cutting site (DEVDA), His- and FLAG tags at the C-terminus^50^. For the structural studies of AKT1-*At*KC1 and AKT1-*At*KC1(G315D), a FLAG tag and a Strep tag were added to the C-terminus of AKT1 and that of the *Arabidopsis thaliana At*KC1 or *At*KC1(G315D), respectively. All the constructs were transiently transfected to and overexpressed in HEK293F cells with a starting density at 2×10^6^ cells ml^−1^ in 5% CO_2_ at 37°C. For AKT1-*At*KC1 or AKT1-*At*KC1(G315D) complex, the two plasmids were transfected as a mixture with a mass ratio of 1:1. The transfected cells were collected by centrifugation (4000g for 10min) 48h after the transfection and stored at −80°C until use.

To purify AKT1, 2L of thawed cell pellets were resuspended in extraction buffer I (25 mM Tris-HCl pH 8.0, 200 mM KCl, 1 mM DTT, 1 μM leupeptin, 1 μM pepstatin A, 1 μg/mL aprotinin, 2 mM PMSF, 0.5% LMNG : CHS (10 : 1, w/w)) and incubated at 4 °C for 2 hours with gentle agitation. Supernatant from centrifugation at 12000 rpm for 45 min was applied to 2 mL of Anti-DYKDDDDK G1 Affinity Resin (GenScript) which was washed five times by 5 mL of wash buffer I (25 mM Tris-HCl pH 8.0, 200 mM KCl, 1 mM DTT, 1 μM leupeptin, 1 μM pepstatin A, 1 μg/mL aprotinin, and 0.02% GDN). The target protein was eluted with 200 μg/mL FLAG peptide in wash buffer I, and subsequently concentrated by Amicon Ultra-4 centrifugal filter (MWCO 100 kDa) before being injected into a Superose 6 Column (GE Healthcare). The size exclusion chromatography (SEC) was performed in wash buffer I and peak fractions were collected and concentrated to 9 mg/ml for cryo-EM sample preparation.

Identical purification protocol was followed to prepare the AKT1-*At*KC1 and AKT1-*At*KC1(G315D) complexes, except that the tandem affinity chromatography was conducted to prepare homogeneous AKT1-*At*KC1 and AKT1-*At*KC1(G315D). Briefly, the co-transfected cell pellets were lysed in extraction buffer I, then target proteins were first purified by Anti-DYKDDDDK G1 Affinity chromatography, followed by Strep-Tactin® chromatography to isolate AKT1-*At*KC1 and AKT1-*At*KC1(G315D). To improve purification efficiency, we incubated the elution from Anti-FLAG G1 resin with the Strep-Tactin® Superflow® resin (IBA Lifesciences) for 3 hours at 4 °C. Then beads were collected by low-speed centrifugation and washed in batch with wash buffer I. The protein was eluted by wash buffer I containing 25mM Desthiobiotin, concentrated and subjected to SEC in a Superose 6 column with wash buffer I. Peak fractions were collected and concentrated to 8-9 mg/ml (Millipore concentrator unit) for cryo-EM sample preparation.

### Cryo-EM sample preparation and data acquisition

Aliquots of 3.5 μL of freshly purified AKT1, AKT1-*At*KC1, and AKT1-*At*KC1(G315D) were placed on glow-discharged 300-mesh holey carbon grids (Quantifoil, Au, R1.2/1.3). The grids were blotted at 8°C and 100% humidity for 2.5 s, 3 s, and 2.5 s, respectively, and rapidly plunged into liquid ethane for vitrification.

The grids were loaded onto a 300-kV Titan Krios (Thermo Fisher Scientific Inc.) equipped with K3 Summit detector (Gatan) and GIF Quantum energy filter. Images were automatically collected using AutoEMation^51^ in super-resolution mode at a nominal magnification of 81,000×, with a slit width of 20 eV on the energy filter. A defocus series ranging from −1.2 μm to −2.2 μm were used. Each stack was exposed for 2.56 s with an exposure time of 0.08 s per frame, resulting in a total of 32 frames per stack and the total dose was approximately 50 e^−^/Å^2^ for each stack. The stacks were motion corrected with MotionCor2^52^ and binned 2 fold, resulting in a pixel size of 1.0773 Å/pixel. Meanwhile, dose weighting was performed^53^. The defocus values were estimated with Gctf^54^.

### Image processing

A diagram for data processing is presented in Fig. S5. A total of 4,150, 2,612, and 4,216 micrographs were collected for AKT1, AKT1-*At*KC1, and AKT1-*At*KC1(G315D), respectively. 2,173,540 particles for AKT1 autopicked by Gautomatch were imported to cryoSPARC^55,56^. 225,687 selected particles from 2D classification were subjected to heterogeneous refinement. Next, local CTF refinement and non-uniform refinement improved the density map to a resolution of 3.12 Å with 145,300 selected particles. 2D classification results of these particles were used as templates for autopicking in Relion^57–59^. 2,462,846 particles, 1,413,304 and 2,603,147 for AKT1, AKT1-*At*KC1 complex, and AKT1-*At*KC1(G315D), respectively, were picked and imported to cryoSPARC. The 3.12 Å map was low-pass filtered as the 3D template. Several rounds of multi-reference heterogeneous classification were carried out on AKT1, AKT1-*At*KC1, and AKT1-*At*KC1(G315D). 399,278, 289,326, and 649,659 selected good particles for AKT1, AKT1-*At*KC1, and AKT1-*At*KC1(G315D), respectively, were subjected to non-uniform refinement without imposing symmetry, resulting in 3D maps with overall resolutions of 2.81 Å, 2.98 Å, and 2.77 Å, respectively. Then, local CTF refinement and non-uniform refinement with C2 symmetry imposed were performed on AKT1 and AKT1-*At*KC1 and yielded density maps at resolutions of 2.69 Å and 2.80 Å, respectively. 2D classification, heterogeneous refinement, local CTF refinement, and non-uniform refinement were performed with cryoSPARC. The resolution was estimated with the gold-standard Fourier shell correlation 0.143 criterion^60^ with high-resolution noise substitution^61^.

### Model building and structure refinement

The coordinates of AlphaFold-predicted structures of AKT1 and *At*KC1 were docked into three maps in CHIMERA^62^. Sequences without clearly resolved densities were deleted in COOT^63^, and G315 of *At*KC1 in AKT-*At*KC1(G315D) was mutated to aspartic acid manually. All three structures were adjusted according to correlative high-resolution maps, and every residue was carefully checked. The chemical properties of amino acids and surrounding environments were considered during model building. In total, 1764, 1830, and 1797 residues for AKT1, AKT-*At*KC1, and AKT-*At*KC1(G315D) were assigned with side chains, respectively.

Structure refinement was performed using phenix.real_space_refine application in PHENIX^64^ in real space with secondary structure and geometry restraints. Over-fitting of the overall model was monitored by refining the model in one of the two independent maps from the gold-standard refinement approach and testing the refined model against the other map^65^. Statistics of the map reconstruction and model refinement can be found in Table S1.

## Data Availability

Atomic coordinates and EM maps of AKT1 (PDB: 7WM2; EMDB: EMD-32598), AKT1-*At*KC1 (PDB: 7WM1; EMDB: EMD-32597), and AKT1-*At*KC1(G315D) (PDB: 9IS8; EMDB: EMD-60834) have been deposited in the Protein Data Bank (http://www.rcsb.org) and the Electron Microscopy Data Bank (https://www.ebi.ac.uk/pdbe/emdb/).

## Acknowledgments

We thank the Cryo-EM Facility and Supercomputer Center of Westlake University for providing data collection and computation support, respectively. We thank Li Huang, Zhipeng Jiang, and Xiaojuan Wang for technical support during cryo-EM image acquisition. We thank Zhenyuan Liu for technical support in computation. This work was supported by the National Natural Science Foundation of China (32122042 and 32071208 to H.S., 82271498 and 82341246 to Z.H.), the STI2030-Major Projects (2021ZD0202103 to Z.H.), the Zhejiang Provincial Natural Science Foundation (DQ24C050001 to H.S.), the Ningxia Hui Autonomous Region Key Research and Development Project (2022BEG02042 to Z.H.), the Westlake Education Foundation (to H.S.), and the Science Foundation of Peking University Cancer Hospital (JC202304 to Z.H.).

## Author information

These authors contributed equally: Zijie Zheng, Yannan Qu, Jiexin Chen.

## Contributions

The project was conceived by H.S. Molecular cloning, protein expression and purification, cryo-sample preparation, and electron micrography data collection were performed by Z.Z., D.L., and Y.T. Y.Q. conducted the structure reconstruction and model building. J.C. carried out electrophysiological experiments under the supervision of Z.H. All authors contributed to the data analysis. H.S wrote the manuscript.

---

**Fig. S1.**
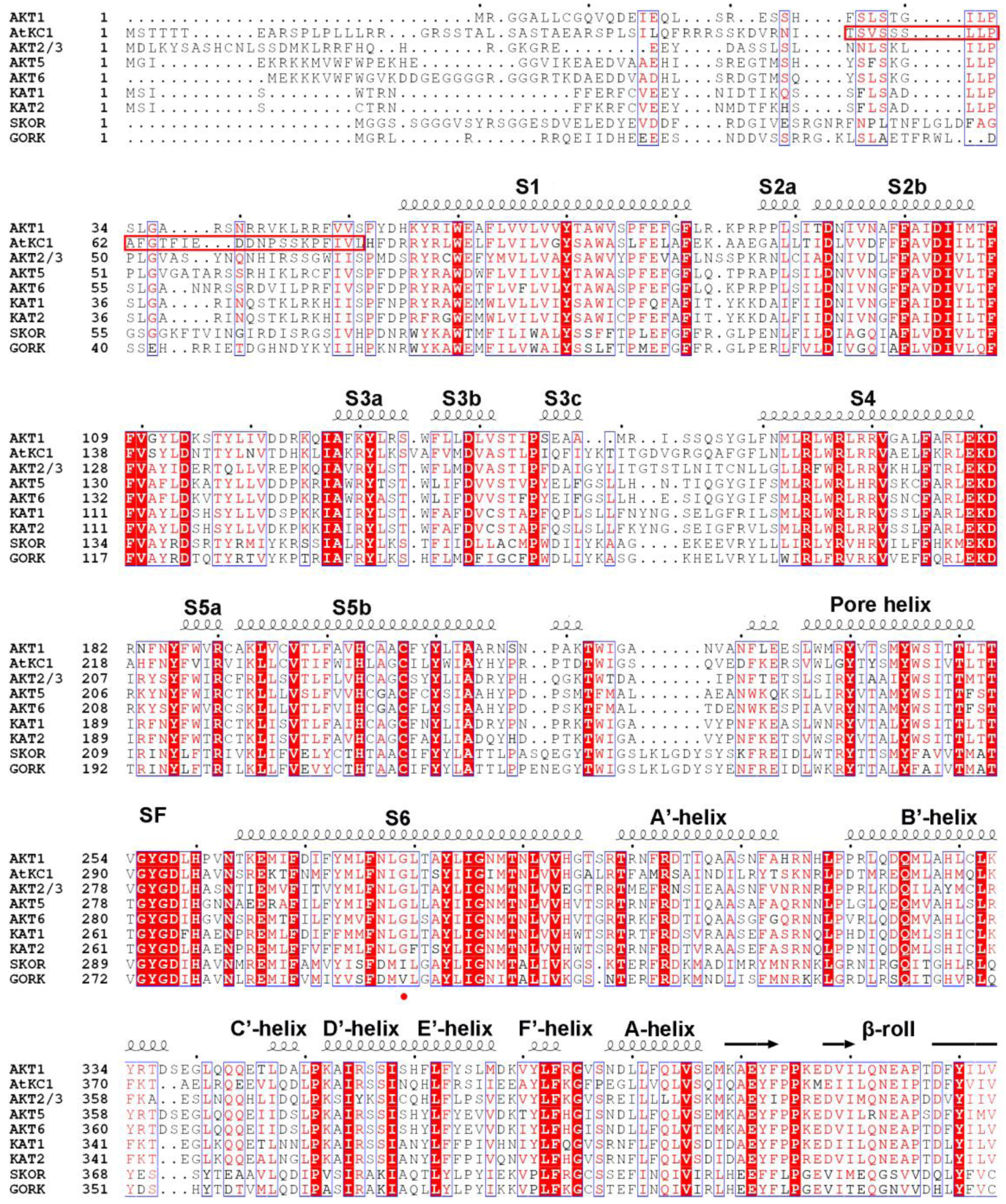
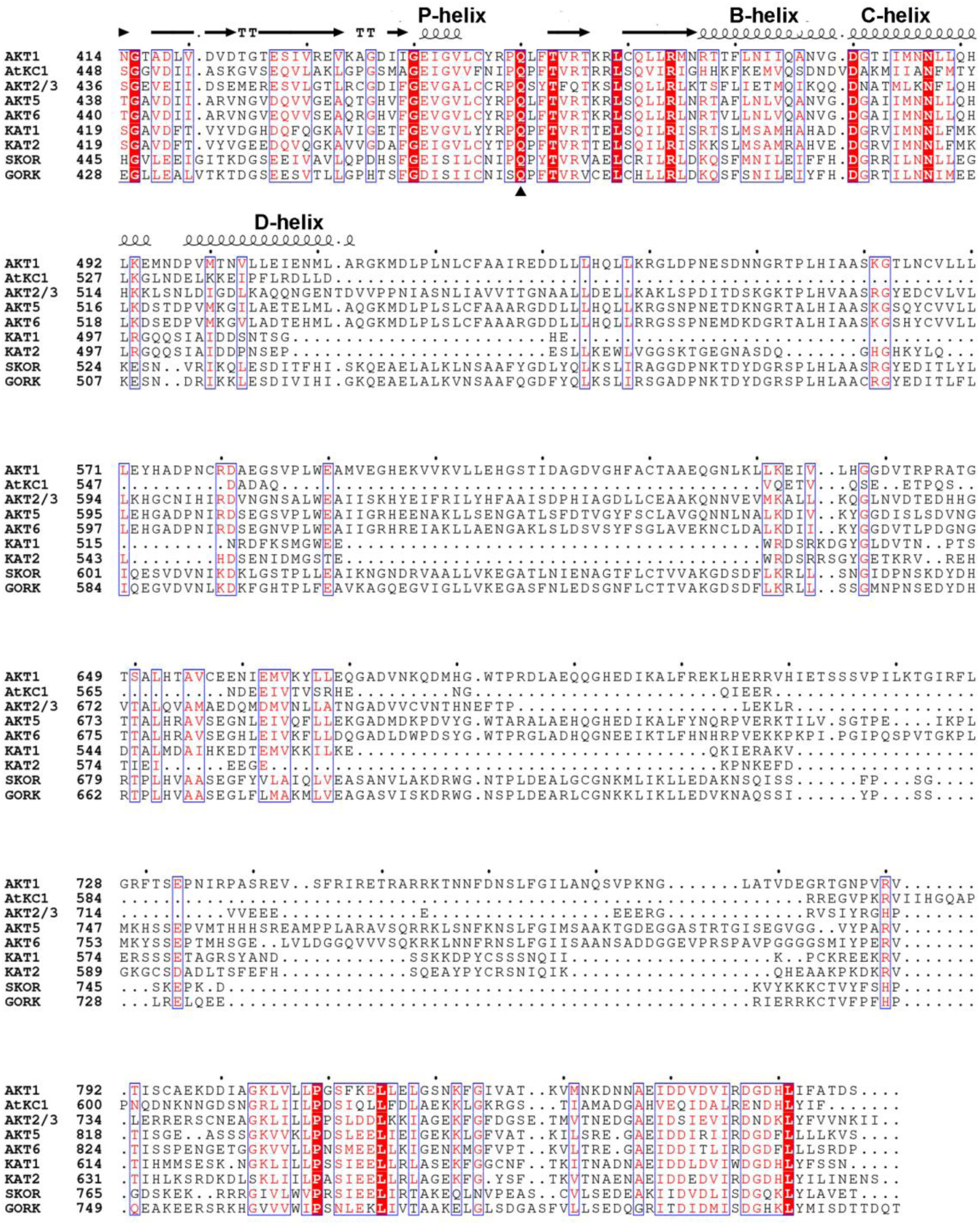
Sequence alignment of *Arabidopsis thaliana* Shaker-like voltage-gated K^+^ channels. The protein sequences of *Arabidopsis thaliana* Shaker-like voltage-gated K^+^ channels are aligned by the Clustal Omega program^68^ and colored with ENDscript 2^69^. The secondary structures of AKT1 are depicted above the aligned sequences. Transmembrane segments, selectivity filter (SF), α helices and β-roll in the C-linker and CNBHD domain are labeled. Invariant residues are shaded in red, while conserved residues are colored red. The I-domain of *At*KC1 (residues 53-80) is highlighted by enclosing it with rectangles, while Gly315 is indicated with a filled red circle. The position of the critical arginine in bona fide CNBD domains is occupied with a glutamine residue, indicated by the filled triangle, in these Shaker-like K^+^ channels, suggesting that these channels are not gated by cyclic nucleotides^37^.

**Fig. S2.**
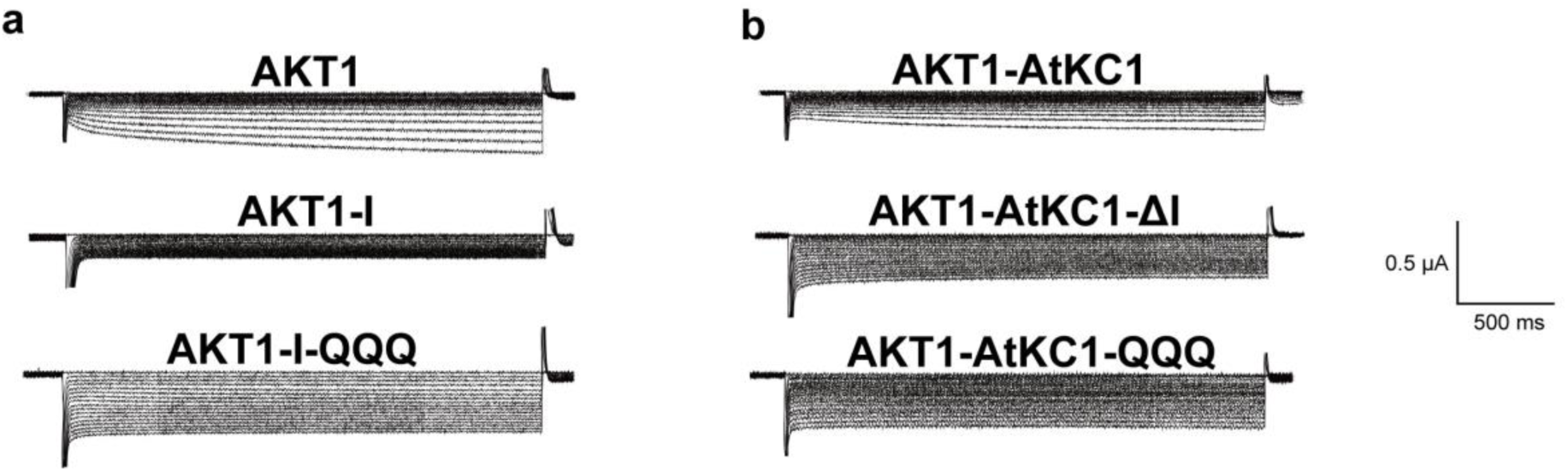
Representative traces of electrophysiological recordings. (**a**)(**b**) Representative traces of AKT1, AKT1-I, AKT1-I-QQQ (**a**), and AKT1-*At*KC1, AKT1-*At*KC1-ΔI, AKT1-*At*KC1-QQQ (**b**) were recorded in bath solutions containing indicated concentrations of K^+^.

**Fig. S3.**
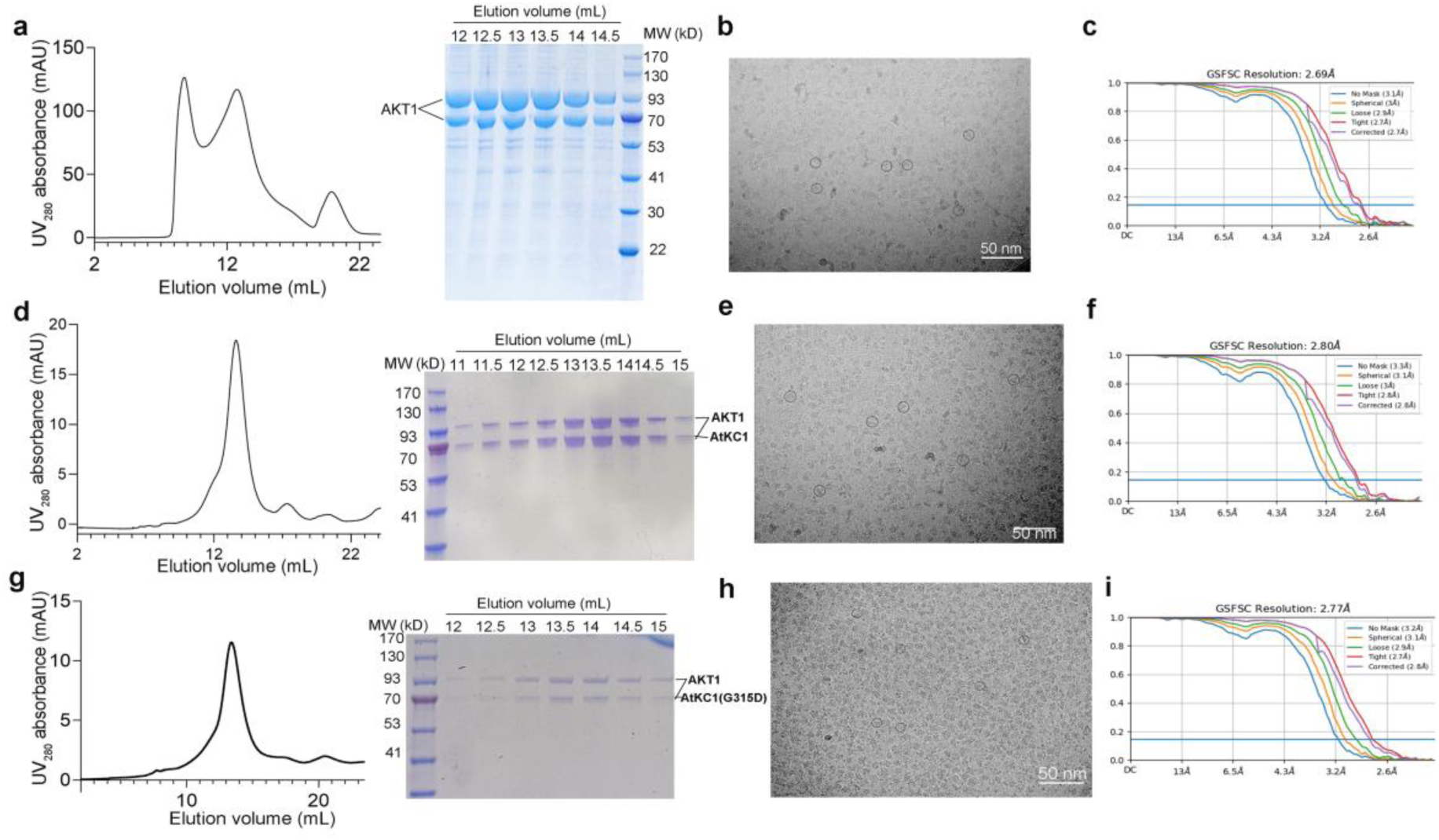
Protein purification and data processing of AKT1, AKT1-*At*KC1, and AKT1-*At*KC1(G315D). (**a**, **d**, **g**) Size exclusion chromatography (SEC) profiles and the correlated Coomassie blue-stained polyacrylamide gel electrophoresis (PAGE) gels of AKT1 (**a**), AKT1-*At*KC1 (**d**), and AKT1-*At*KC1(G315D) (**g**). (**b**, **e**, **h**) Protein purification details can be found in Materials and methods. Representative micrographs of AKT1 (**b**), AKT1-*At*KC1 (**e**), and AKT1-*At*KC1(G315D) (**h**) are shown. (**c**, **f**, **i**) FSC curves for the final round of cryo-EM map reconstructions of AKT1 (**c**), AKT1-*At*KC1 (**f**), and AKT1-*At*KC1(G315D) (**i**) are displayed, indicating that the resolutions of the final reconstruction are 2.69, 2.80, and 2.77 Å, respectively, according to the 0.143 cutoff criterion^60^.

**Fig. S4.**
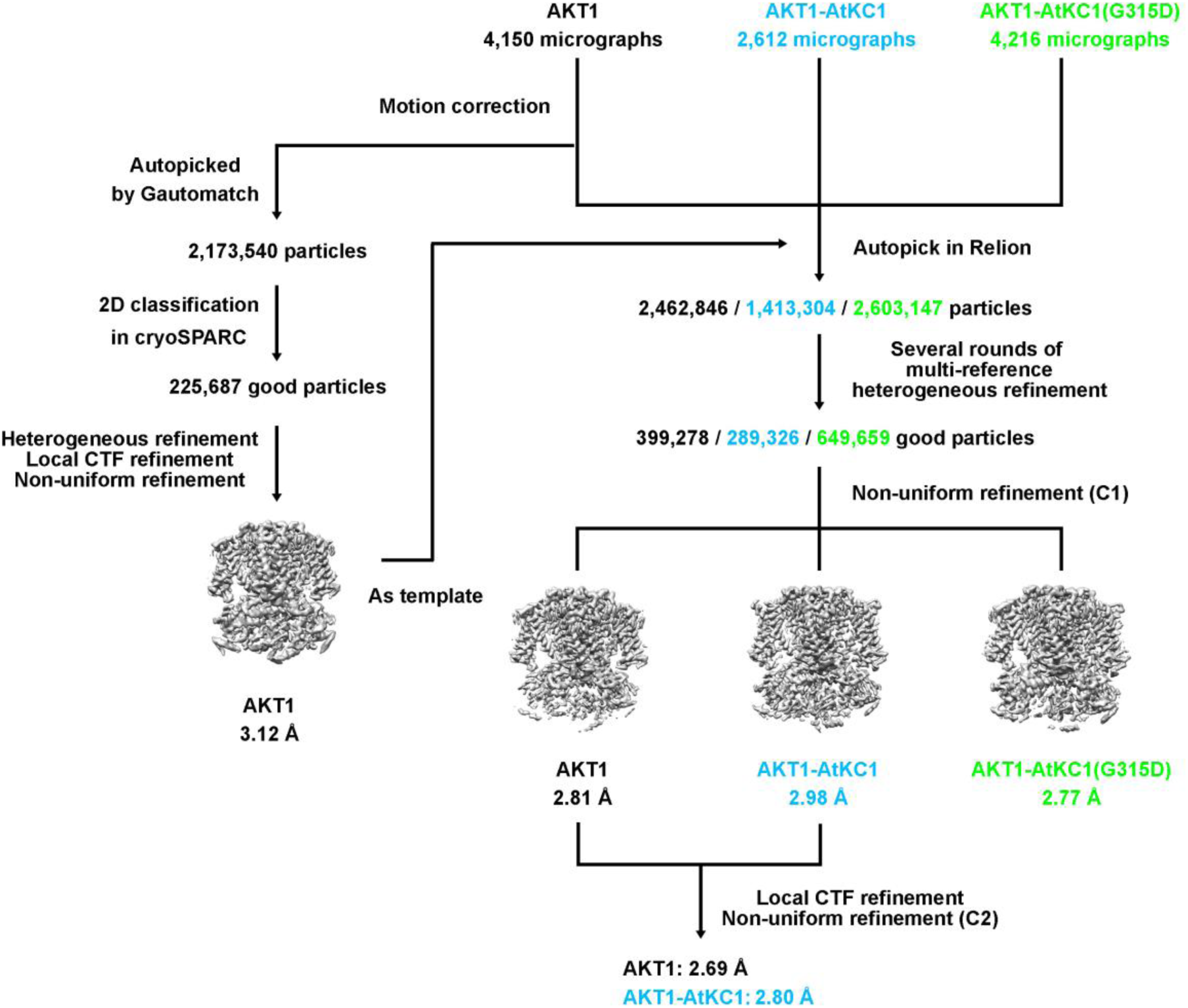
Flowchart of EM data processing for AKT1, AKT1-*At*KC1, and AKT1-*At*KC1(G315D). EM maps of AKT1, AKT1-*At*KC1, and AKT1-*At*KC1(G315D) were reconstructed at final resolutions of 2.69 Å, 2.8 Å, and 2.77 Å, respectively. For details on cryo-EM data processing, please refer to the “Image processing” section in Materials and methods.

**Fig. S5.**
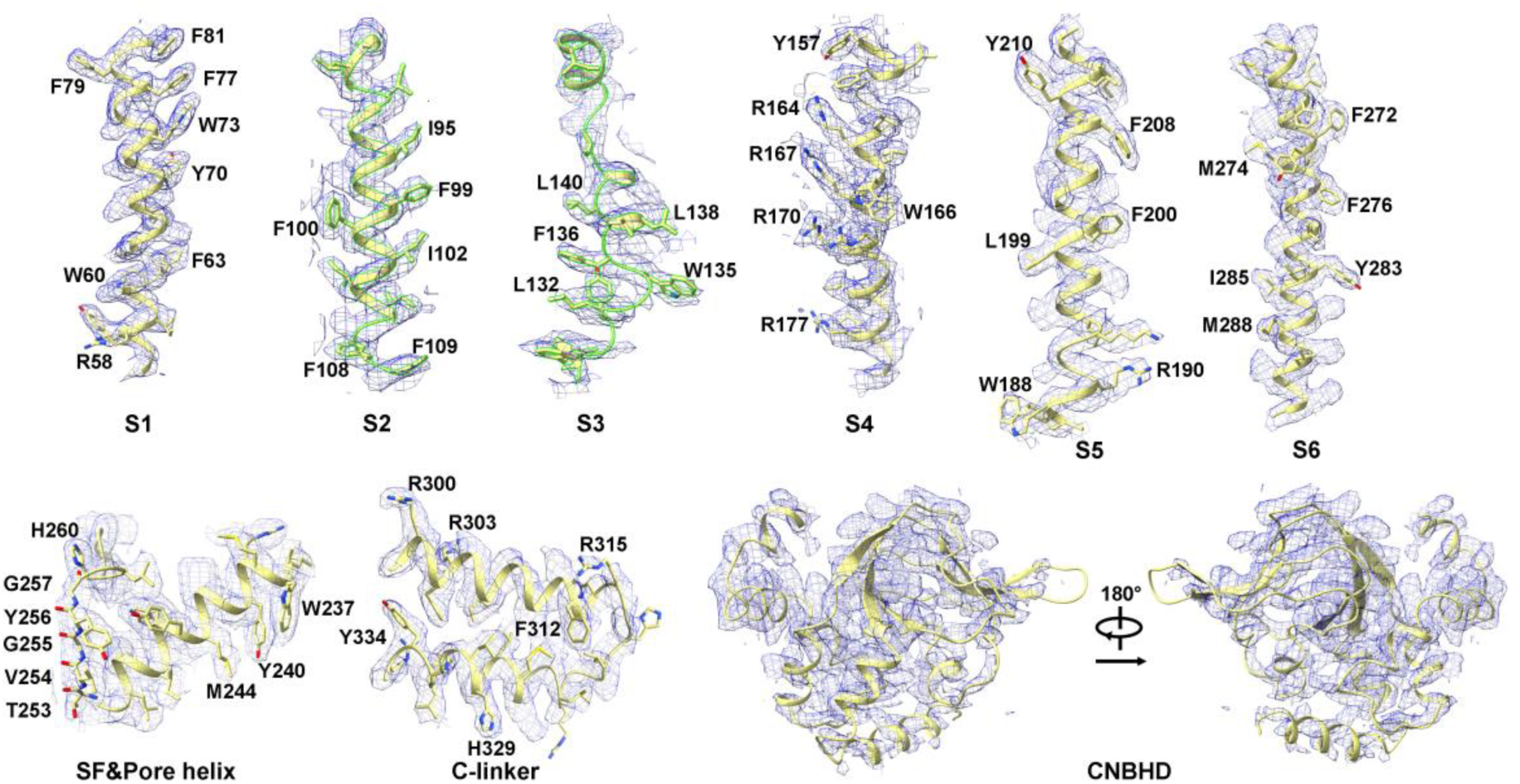
Representative EM densities for the “kinked” AKT1 in AKT1-*At*KC1(G315D). Representative densities for the “kinked” AKT1 in AKT1-*At*KC1(G315D) are displayed. The figures were prepared in UCSF ChimeraX^66^.

**Fig. S6.**
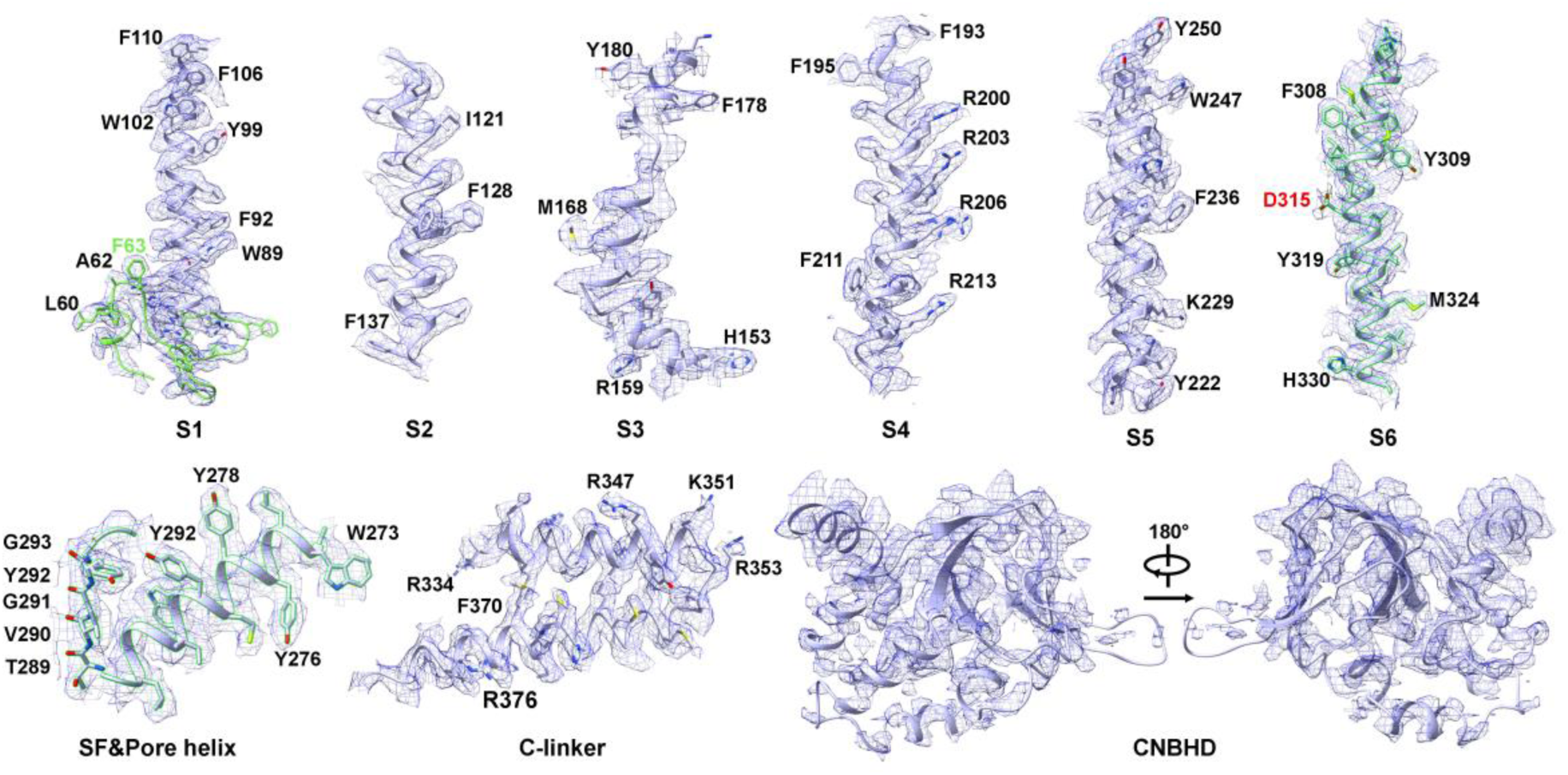
Representative EM densities for *At*KC1(G315D). Representative densities for *At*KC1(G315D) in AKT1-*At*KC1(G315D) are displayed. The I-domain is shown in green and residue Asp315 is highlighted in red. The figures were prepared in UCSF ChimeraX^66^.

**Fig. S7.**
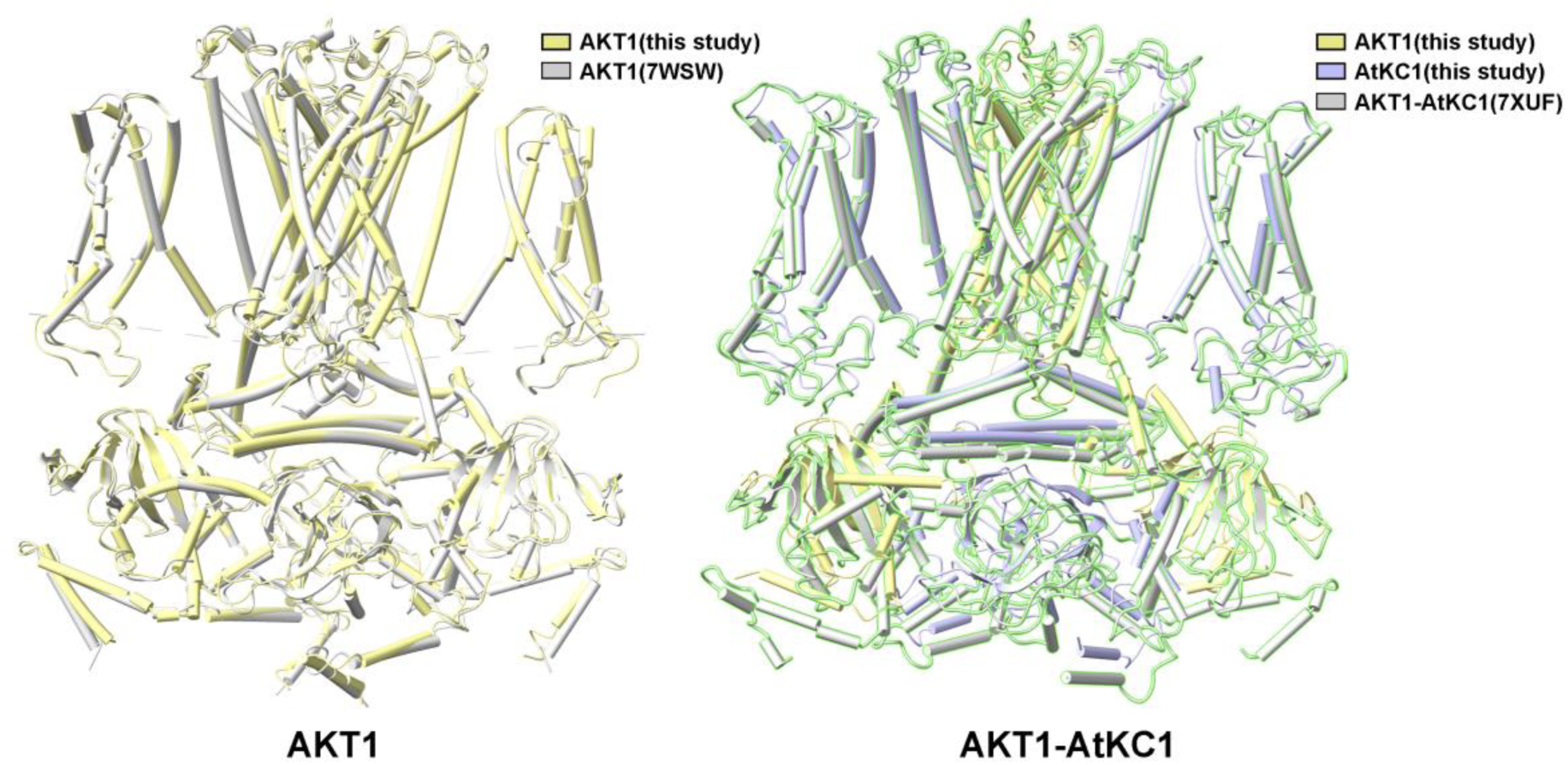
Alignment between our structures with previously reported ones. AKT1 (*left*) and AKT1-*At*KC1 (*right*) structures determined in this study and reported previously by other groups are aligned and compared using UCSF ChimeraX. The alignment reveals a high similarity between our structures and the ones reported previously. AKT1 (PDB code: 7WSW) and AKT1-*At*KC1 (PDB code: 7XUF) structures are used for structural comparison.

**Fig. S8.**
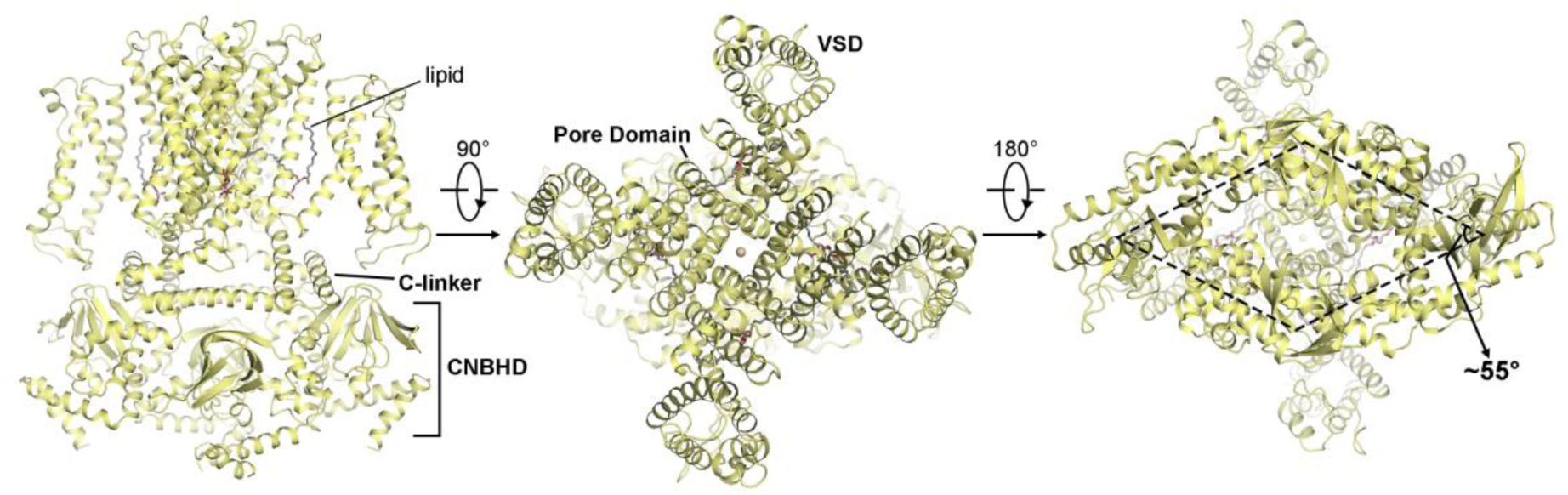
Overall structure of AKT1. Side, top, and bottom views of AKT1 are illustrated. Individual domains, including the C-linker, CNBHD, Pore, and Voltage-sensing (VSD) domains of AKT1 and the lipids intercalated between the pore domain and VSD are labeled. The C-linker and CNBHD domains of AKT1 exhibit a rhombus shape with an interior angle of about 55°.

**Fig. S9.**
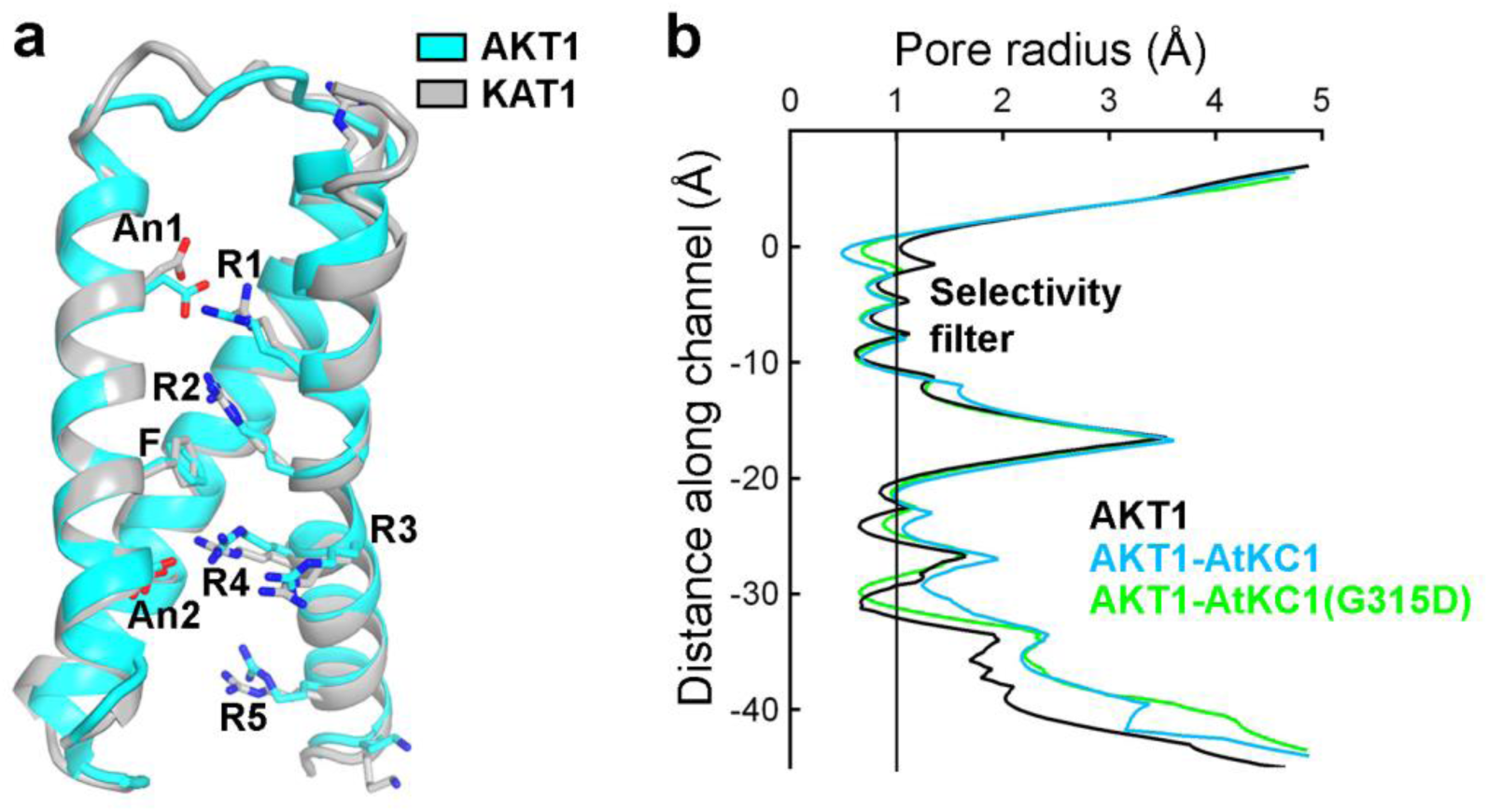
AKT1, AKT1-*At*KC1, and AKT1-*At*KC1(G315D) structures adopt depolarized, closed conformation. (**a**) The voltage-sensing domain (VSD) of AKT1 is superposed with the depolarized VSD of the KAT1 structure (PDB code: 6V1X)^35^. Both Arg164(R1) and Arg167(R2) are above the hydrophobic constriction site (HCS, Phe100 in AKT1), suggesting that the VSD of AKT1 is also in a depolarized conformation. (**b**) The radii along the permeation pore of AKT1, AKT1-*At*KC1, and AKT1-*At*KC1(G315D), calculated with HOLE, are less than 1 Å, indicating the channels are in closed conformations^70^.

**Fig. S10.**
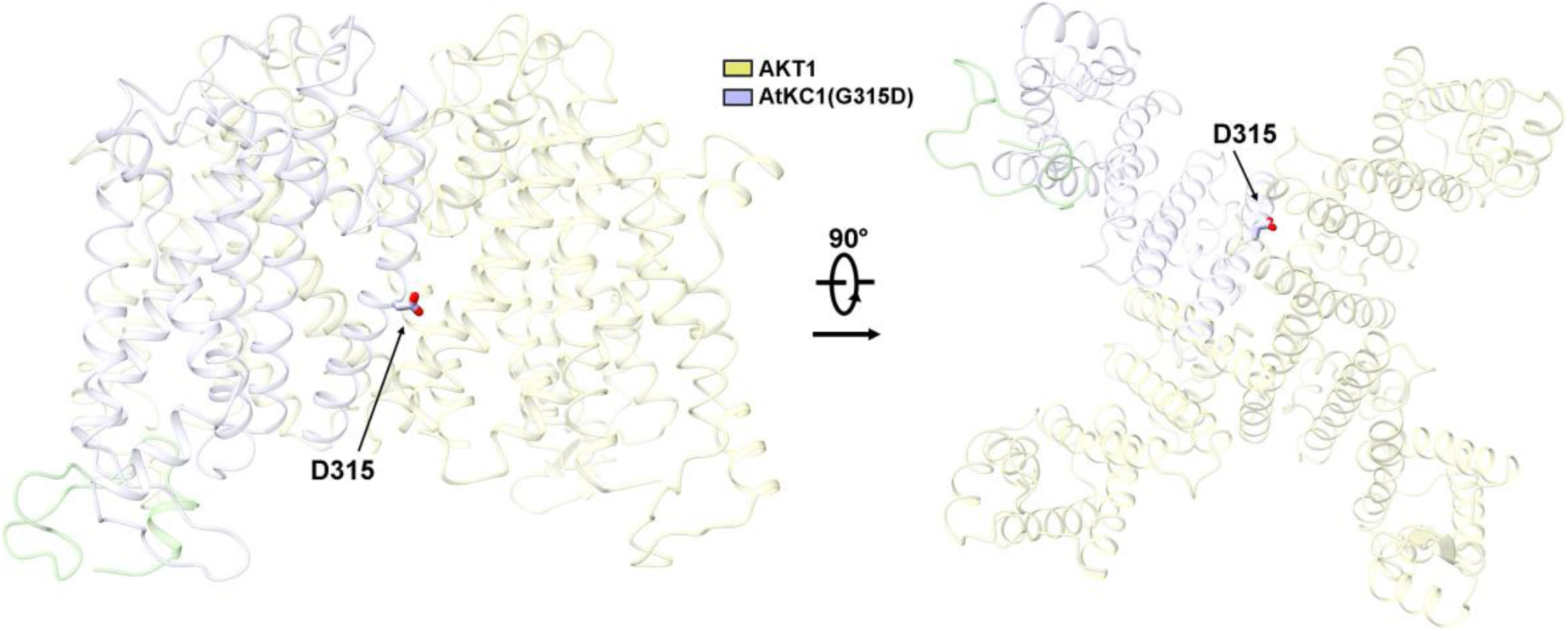
The position of Asp315 in AKT1-*At*KC1(G315D). The side chain of Asp315 in AKT1-*At*KC1(G315D) points unfavorably towards the hydrophobic lipid bilayer in the middle of plasma membranes. For the density of Asp315 in the complex, please refer to Fig. S6.

**Table S1.**
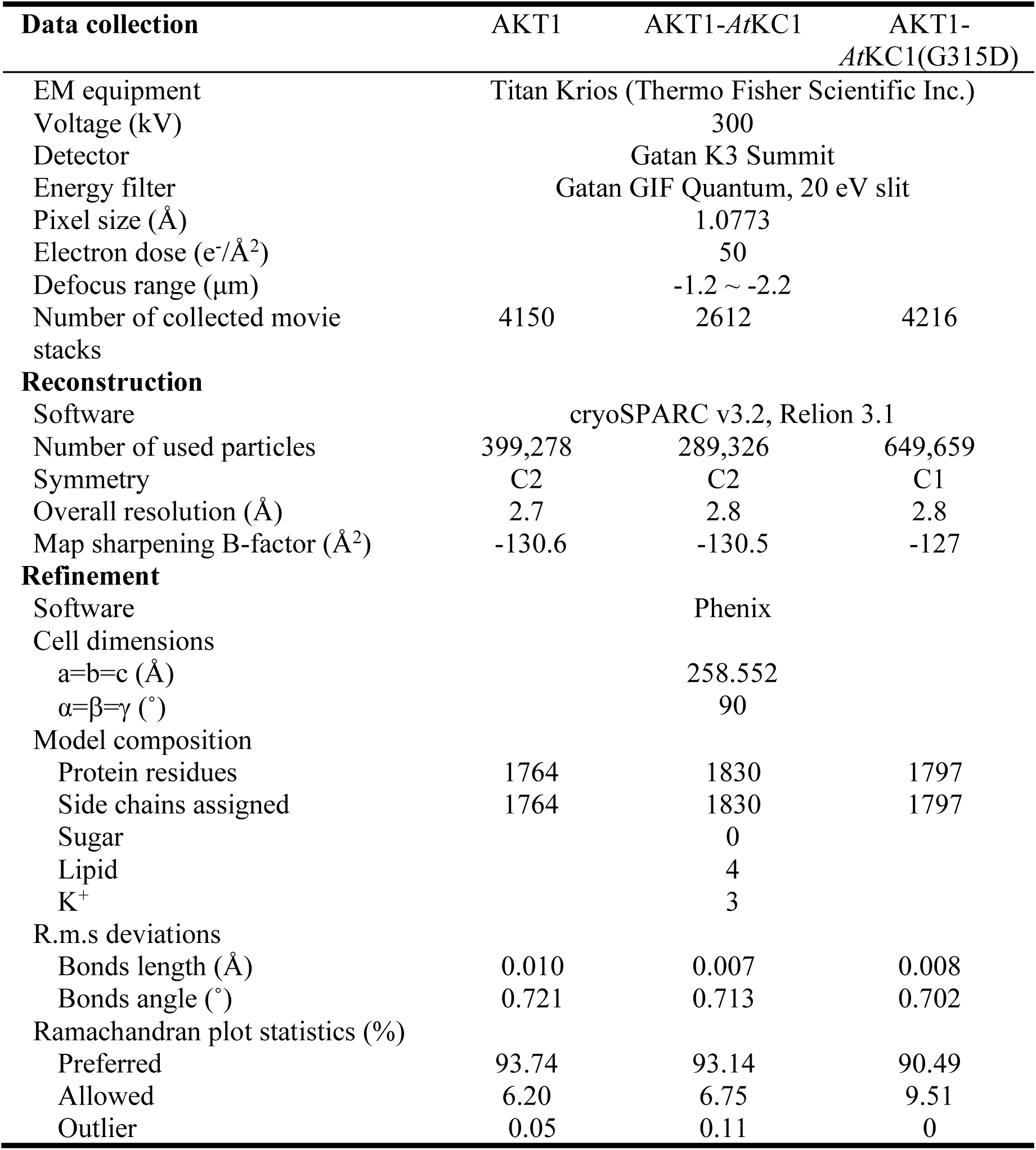
Statistics for data collection and structural refinement.

